# Heavy Metal and Antibiotic Co-resistance in Bacteria Isolated from Poultry Samples in Bangladesh: An Emerging Environmental Threat

**DOI:** 10.1101/2024.10.25.620196

**Authors:** Amily Sarker, Nabila Haque, Mohammad Ayaz, Umme Samia Antu, Taslin Jahan Mou, Shuvra Kanti Dey, Md.Anowar Khasru Parvez

## Abstract

The excessive use of heavy metals and antibiotics as additives in livestock feed may significantly contribute to bacterial resistance in poultry environments by exerting selective pressure. This study aimed to comprehensively analyze bacterial isolates from poultry farms across Bangladesh, focusing on their resistance to heavy metals and antibiotics. Through microbial phenotypic screening, 23 cadmium-resistant and 28 chromium-resistant bacterial isolates were identified. The minimum inhibitory concentrations (MICs) for cadmium and chromium in metal-resistant bacterial isolates varied significantly, ranging from 200 μg/ml to 1600 μg/ml. All cadmium-resistant isolates were fully resistant to cefixime and ceftazidime, while chromium-resistant isolates showed 90% resistance to tetracycline. The cadmium resistance gene, such as *czc*, was detected in 26% (6/23) of cadmium-resistant isolates, and the chromium reductase gene *chrR* was detected in 18% (5/28) of chromium-resistant isolates. Among the chromium-reductase isolates, 40% (2/5) exhibited co-resistance and harbored the *chrR* gene, beta-lactam resistance genes (*bla-TEM*, *bla-NDM*), and the colistin resistance gene (*mcr-2*). A significant association (p = 0.026) was observed between the presence of the *chrR* gene and these antibiotic resistance genes. However, no co-resistance was found in cadmium-resistant isolates. Mobile genetic elements, specifically Class 1 integrons, were found in 33.33% (2/6) of the isolates carrying the cadmium resistance gene (*czc*) and in 100% (5/5) of the isolates containing the chromium reductase gene (*chrR*). These findings underscore poultry environments in Bangladesh as significant reservoirs for bacteria with dual resistance to heavy metals and antibiotics, emphasizing the need for further investigation into the genetic mechanisms driving microbial resistance and its implications for public health.

## Introduction

Intensive poultry farming has become highly efficient, evolving to meet animal dietary needs and increasing consumer demand globally. Nevertheless, such high production levels are linked to a considerable environmental impact, primarily due to waste by-products that affect the interconnectedness of animals, their surrounding environment, and humans [1,2]. Elements like copper (Cu), zinc (Zn) from feed supplements, extensively used in the chicken industry for therapeutic and growth purposes, have been found in chicken tissues [3]. These heavy metals, due to their durability and persistence, can lead to the emergence of metal resistance in bacteria within food and livestock, potentially causing environmental contamination from the residual metals on poultry farms [4].

The growing antimicrobial resistance is now widely seen as a serious global health challenge, increasing risks to both human and animal health around the world [5]. The emergence of bacteria producing β-lactamase (*blaTEM, blaSHV, blaNDM-1,ctxM15*), carbapenems resistant (*blaKPC*), and colistin-resistant (*mcr-1, mcr-2*) has been a concern in both veterinary and human medicine due to their potential resistance to all available antimicrobials and their association with high morbidity and mortality rates in poultry [6–9]. The selective pressure from heavy metals is recognized as a significant factor in spreading antibiotic resistance genes (ARGs), with livestock waste as a source of antibiotics and heavy metals [10]. This results in the simultaneous contamination of farms and their surroundings with antibiotics and metals, thereby raising the chances of bacteria acquiring antibiotic resistance genes (ARGs) and promoting the spread of ARG contamination in the environment [11].

Recent research has revealed a direct link between heavy metals and antibiotic resistance genes (ARGs), indicating that heavy metals contribute to their increased prevalence by co-selecting for them [12,13]. Metal resistance genes (MRGs) and antibiotic resistance genes (ARGs) are co-selected via multiple environmental mechanisms. [14]. Mobile genetic elements, including insertion sequences, integrons, and transposons, play a vital role in developing microbiomes containing antibiotic resistance genes (ARGs), as they enable horizontal gene transfer between microorganisms [15–18]. Class 1 integrase genes (*intI-1*) are prevalent in bacteria from poultry litter, regardless of antibiotic use, enabling the capture of various ARG cassettes by class 1 integrons and the propagation of these cassettes by transposons [15,17].ARGs and MRGs are understood to have similar mechanisms of action, with environmental metal stress acting as a co-selection agent that facilitates the spread of ARGs, leading to environmental pollution. The genetic exchange within bacterial communities under environmental stress strengthens bacterial defense mechanisms against these stressors [19]. Recent research has focused on ARG contamination in the farming environment, particularly in poultry feces [20], cloacal swabs [21], and nearby farm sewage [22]. However, studies on the co-occurrence of heavy metal and antibiotic resistance in bacteria from Bangladeshi poultry farms are scarce. This study aims to investigate bacteria’s resistance to heavy metals, both phenotypically and genotypically, understand the relationship between heavy metal resistance genes (HMRGs) and ARGs, and identify the presence of mobile genetic elements in transmitting resistance.

## Materials and Methods

### Sampling, Transportation, and Physicochemical Analysis

The study, conducted between June and December 2022, covered seven districts across five divisions of Bangladesh: Savar, Manikgonj, Tangail (Dhaka division), Natore (Rajshahi), Noakhali (Chittagong), Gaibandha (Rangpur), and Barisal (Barisal division). These regions were selected due to their large poultry populations and frequent disease reports. Samples were gathered from seven distinct locations throughout Bangladesh, including seven poultry droppings samples from farms and two surface water samples from nearby locations. The droppings were placed in sterile falcon tubes. At the same time, water samples were stored in 500 ml sterile Duran bottles (Schott, Germany), securely sealed and transported to the lab in insulated ice boxes for immediate bacteriological analysis. Physicochemical parameters such as temperature, pH, and dissolved oxygen (DO) were measured using a thermometer, pH meter electrode (Orion-2 STAR, Thermoscientific), and a DO meter (970 DO2 meter, Jenway UK).

### Primary screening for Cadmium and Chromium-resistant bacteria

The screening of cadmium and chromium-resistant bacteria was carried out using nutrient agar (HIMEDIA, India) plates containing 100 µg/ml of Potassium dichromate [K_2_Cr_2_O_7_] (Loba Chemie, India) and Cadmium Sulphate Hydrate[3CdSO_4_.8H_2_O] (Loba Chemie, India) in the medium [23]. Nutrient agar media supplemented with chromium and cadmium were prepared using distilled water, with the pH adjusted to 7.5. First, 5g of each poultry droppings was diluted in 5 mL sterile distilled water. The samples were ten-fold diluted for isolation of bacteria, ranging from 10^-1^ to 10^-5^. A 100µl sample from each dilution was aseptically transferred using a micropipette to individual nutrient agar plates supplemented with cadmium and chromium. The plates were incubated for 24 hours at 30°C.Following incubation, the plates were examined for distinct bacterial colonies exhibiting variations in morphology, including shape, elevation, margin, surface characteristics, and size. The total bacterial count was measured in terms of colony-forming units (cfu) per milliliter. Single colonies were picked out and isolated through streak plate cultivation to get pure culture. The purified and grown well colonies were stored under −20 °C.

### Biochemical characterization and identification of representative colony

Representative isolates were then subjected to biochemical tests for characterization, including indole production, methyl-red and Voges-Proskauer test, citrate utilization test, TSI agar test and catalase production. Each test was accompanied by (+) ve and (-) ve controls. The biochemical results were summarized using the online tools Microrao: http://www.microrao.com/index.html and Bio Cluster. This platform predicts potential organisms by analyzing biochemical data [24].

### Determination of the minimum inhibitory concentrations (MICs) of heavy metals

There is no established standard procedure for assessing bacterial susceptibility to heavy metals. The lowest concentration of heavy metals required to inhibit bacterial growth (MIC) was measured for metal-resistant strains using nutrient agar (HIMEDIA, India) supplemented with chromium and cadmium at concentrations ranging from 100 to 3200 µg/mL [23, 25]. Solutions of metal salts were first prepared in sterilized deionized water and then added to MH agar at varying concentrations. Approximately 3 × 10⁶ cells were inoculated onto the agar and then incubated for 18–24 hours at 30°C [25].

### Multiple metal resistance capacity

All cadmium and chromium-resistant isolates were tested for their ability to determine their resistance to other heavy metals such as nickel and lead. Each isolate was individually cultivated on NA agar plates supplemented with cadmium (Cadmium Sulphate, 3CdSO_4_.8H_2_O) (Loba Chemie, India), Chromium (Potassium dichromate K_2_Cr_2_O_7_) (Loba Chemie, India), nickel (Nickel Chloride Hexahydrate, NiCl_2_.6H_2_O) (Loba Chemie, India), and lead (Lead Nitrate, Pb(NO_3_)_2_) (Merck Chemicals, India) at a concentration of 100µg/mL, under pH 7.0 and at 30°C for 24 hours. Following incubation, the isolates were evaluated for their resistance to multiple heavy metals [23].

### Antibiotic Susceptibility Testing

The antibiotic susceptibility of metal-resistant isolates to 16 different antibiotics was tested using the disk diffusion method on Mueller Hinton (MH) agar (HIMEDIA, India), adhering to the protocols outlined by the Clinical and Laboratory Standards Institute (CLSI) in 2011 [26]. The 16 antibiotic disks (obtained from Titan Media Ltd, India) represented eight different classes and included Macrolides (azithromycin 15 mcg); Penicillin (ampicillin ten mcg); Aminoglycosides (gentamycin ten mcg, amikacin 30 mcg); Chloramphenicol (chloramphenicol 30 mcg); Cephalosporins (cefixime 5 mcg, cefuroxime 30 mcg, ceftriaxone 30 mcg, ceftazidime 30 mcg, cefotaxime 30 mcg); Quinolones (ciprofloxacin 5 mcg, levofloxacin 5 mcg, and nalidixic acid 30 mcg); Carbapenem (meropenem 10 mcg); and Tetracycline (tetracycline 30 mcg). Fresh cultures were inoculated into Luria Bertani (LB) broth (HIMEDIA, India) and incubated until turbidity equivalent to a 0.5 McFarland standard was reached. Subsequently, bacterial cultures were plated onto MH agar (HIMEDIA, India) plates in a sterile environment, followed by antibiotic disks. The plates, after inoculation, were incubated for 24 hours at 30°C and the inhibition zone diameters were measured to categorize the isolates as resistant, intermediate or susceptible in accordance with the 2011 CLSI guidelines [26,27]. Isolates that resisted three or more antibiotic classes were categorized as multidrug-resistant (MDR). Furthermore, the multiple antibiotic resistance (MAR) index was calculated using the formula: MAR index = (percentage of antibiotics to which the isolate shows resistance) / (total number of antibiotics exposed to the isolate) [28].

### Molecular characterization of cadmium resistance and chromium reductase gene

PCR was performed to identify resistance gene patterns. The reaction mixture comprised 25 μl volume, including 12.5 μl of master mix, one microliter of both forward (10 pmol) and reverse primers (10 pmol), 6.5 μl of nuclease-free water, and four μl of DNA template. For the detection of cadmium resistance *czc* gene, the following thermal cycling conditions were applied: initial denaturation of template DNA at 95°C for 5 min, followed by 36 cycles of denaturation at 94°C for 1 min, annealing at 61.6°C for 30 s, extension at 72°C for 1 min, and a final extension at 72°C for 10 min. The cooling temperature was maintained at 4°C.

For amplification of the chromium reductase *chrR* gene, PCR was carried out at 95°C for 5 min, followed by 35 cycles consisting of denaturation at 95°C for 30 s, annealing at 55.5°C for 30 s, extension at 72°C for 1 min, and a final extension at 72°C for 10 min. The resulting amplified products were then separated by agarose gel electrophoresis on a 1% gel stained with ethidium bromide and examined under a UV transilluminator.

### Detection of antibiotic resistance gene among metal resistance determinants

The co-existence of metal-resistant isolates was checked for antibiotic resistance determinants using specific primers of variant genes. PCR was utilized to explore the patterns of antibiotic resistance genes. Genes responsible for Class A β-lactamase (*blaTEM, blaSHV, bla-ctxM15, blaNDM-1*), carbapenemase (*blaKPC*) and colistin resistance (*mcr-1, mcr-2*) were detected through PCR using specific primers. These primers were used to generate PCR products, the sizes of which are detailed in **Table 1**.

**Table 1.**
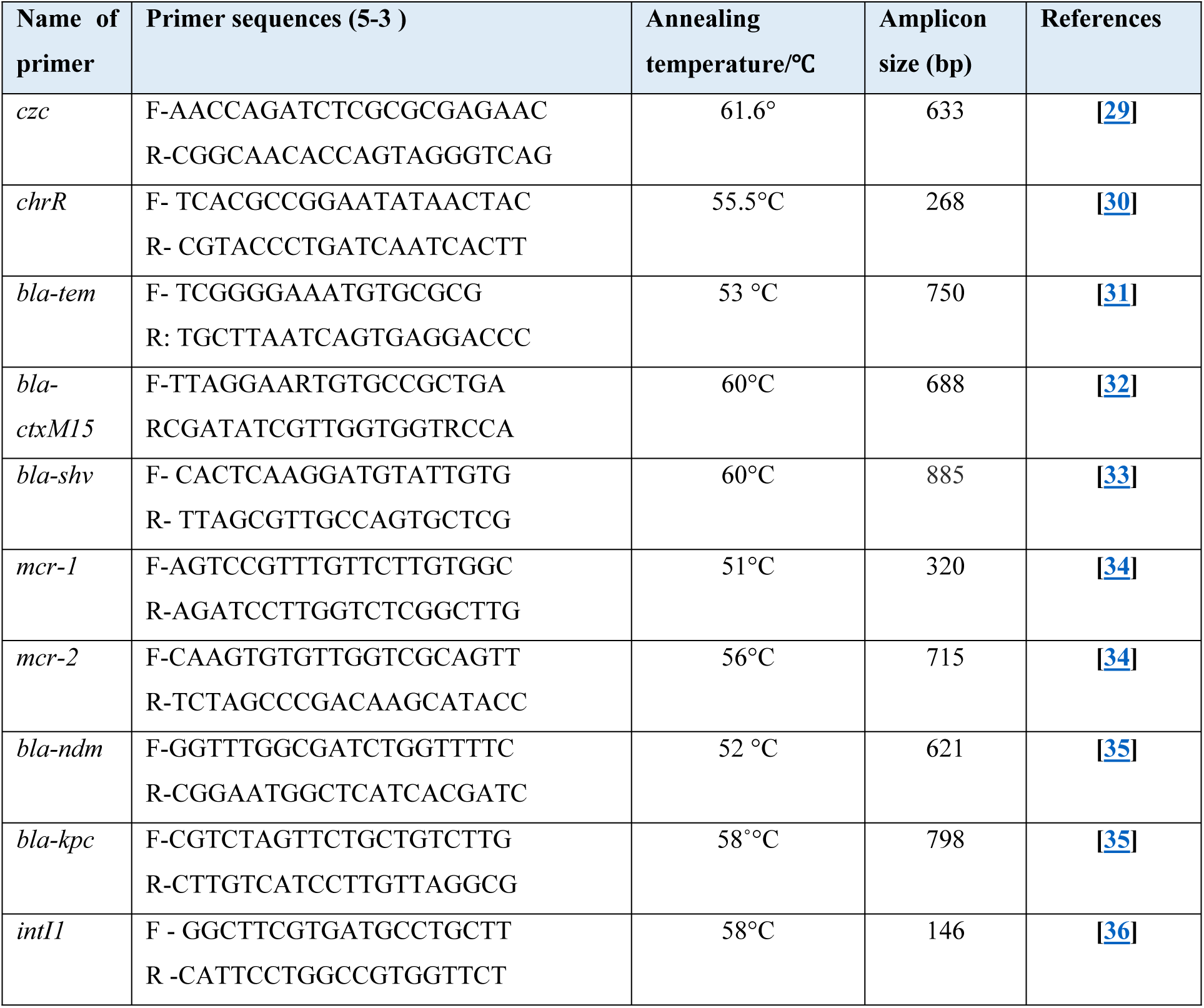
Primer sequences used for detection of bacterial heavy metal resistance and antibiotic resistance and corresponding annealing temperature used for PCR.

### Detection of Class 1 Integrase gene

Integrase genes intI1 were detected in metal-resistant isolates via PCR, using specific primers detailed in Table 1. The PCR process involved a reaction mixture of 25 µL, following previously described protocols, with the reaction conditions including an initial pre-incubation at 94°C for 1 minute, then 35 cycles of 98°C for 30 seconds for denaturation, 58°C for 30 seconds for annealing, and a final elongation phase was conducted for 10 minutes at 72°C. Following amplification, the PCR products were subjected to electrophoresis on a 1% agarose gel, as detailed in **Table 1**.

## Results

### Analysis of physicochemical parameters and Isolation of metal-resistant bacteria from samples

Seven poultry-dropping and two surface water samples were collected to analyze physicochemical parameters and isolate metal-resistant bacteria. The temperature ranged from 24°C to 29.4°C, pH from 5.7 to 9.3, and dissolved oxygen (DO) levels from 4 mg/L to 5.6 mg/L.

Heavy metal-resistant bacteria were isolated using nutrient agar with 100 μg/mL metal. Out of 51 isolates, 28 were resistant to chromium and 23 to cadmium, selected for further study based on colony morphology, size, and color.

### Morphological and Biochemical Characterization

The phenotypic characteristics of these organisms and their tentative identification based on biochemical tests (Gram staining, indole production, methyl red, Voges Proskauer test, citrate utilization test, nitrate reduction test, and Kligler Iron Agar test) were performed. Based on these characteristics, four genera of bacteria, such as *Enterococcus spp., Escherichia coli*, *Staphylococcus spp.*, and *Bacillus spp.*, were identified.

### Minimum inhibitory concentrations (MIC) of heavy metals for metal-resistant isolates

All 23 cadmium-resistant isolates were tested for Minimum Inhibitory Concentration (MIC) using cadmium concentration at 200 μg/ml to 1000 μg/ml, with 60.86% tolerating 200 μg/ml and 30.43% resisting concentrations above 1000 μg/ml. Similarly, the 28 chromium-resistant isolates were evaluated at chromium concentrations ranging from 200 μg/ml to 3200 μg/ml; all tolerated 200 μg/ml, but only 21.42% showed resistance at 3200 μg/ml. **(Fig 1)**

**Fig 1:**
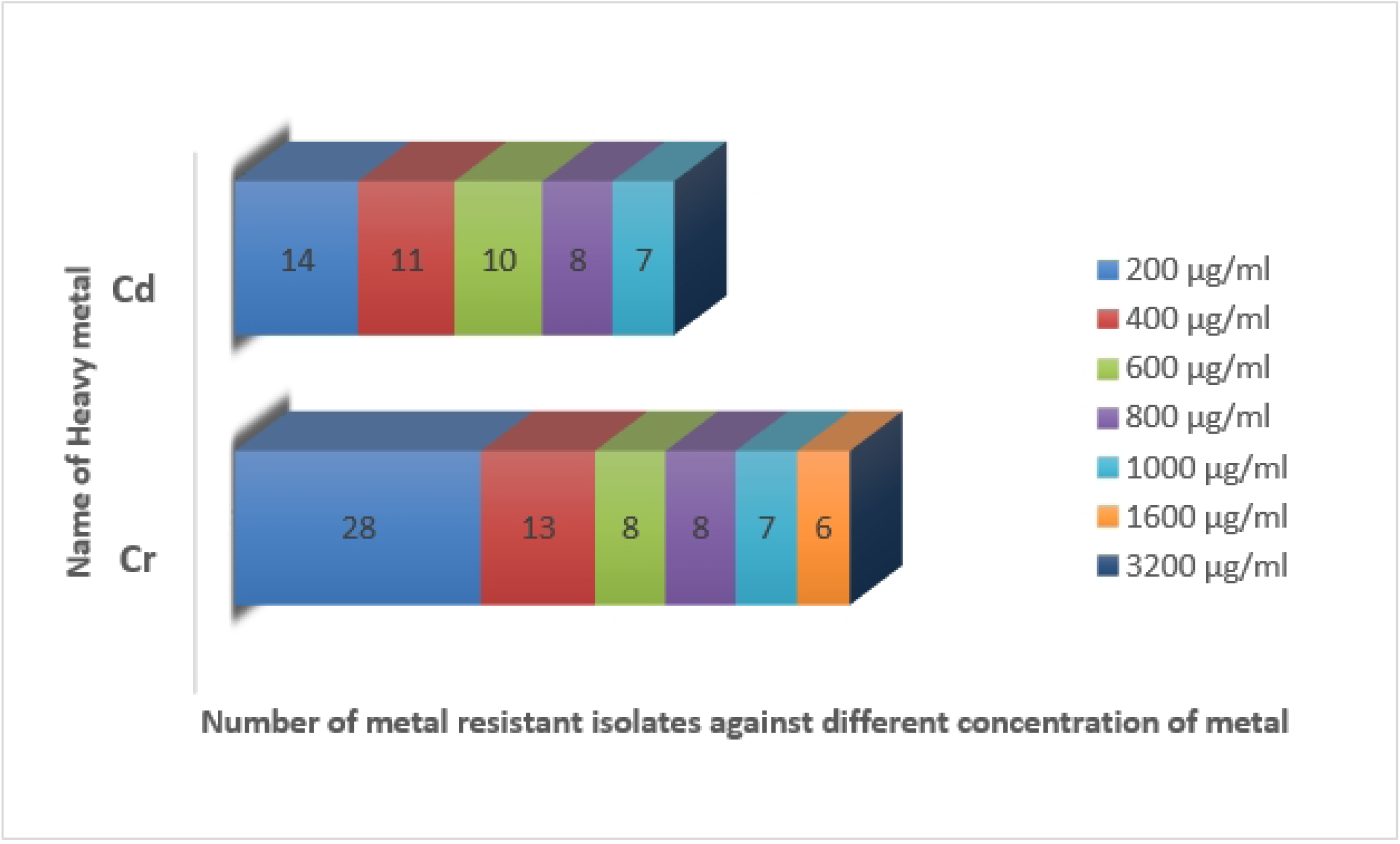
Minimum inhibitory concentrations (MIC) of heavy metals (Cadmium and Chromium) for metal-resistant isolates.

### Determination of Multiple metal resistance capacity

The study observed distinct patterns of Multi Heavy Metal Resistance (MHMR) in bacterial isolates from poultry environments. Among the 23 isolates resistant to cadmium, 44% also withstood lead, 26% showed resistance to nickel, and 52% were resistant to at least two heavy metals. Notably, these cadmium-resistant isolates did not exhibit tolerance to chromium.

Conversely, of the 28 isolates resistant to chromium, 65% demonstrated resistance to lead, 50% to nickel, and 43% were resistant to at least two heavy metals. These chromium-resistant isolates did not show resistance to cadmium. Remarkably, 52% of cadmium-resistant and 79% of chromium-resistant isolates were classified as MHMR, highlighting the diverse and complex nature of heavy metal resistance in these bacteria.

### Antibiotic Susceptibility Profiles of heavy metal-resistant Isolates

The antibiotic resistance profiles of heavy metal-resistant isolates were assessed against 16 antibiotics from eight classes, revealing significant variations across sampling sites. Cadmium-resistant isolates exhibited high resistance to Cefixime and Ceftazidime (100% of isolates), substantial resistance to Tetracycline (63.63%), Nalidixic Acid, and Chloramphenicol (54.54%), moderate resistance to Ampicillin and Levofloxacin (45.45%), and lower resistance to Amikacin (36.36%), Gentamycin (18.18%), with minimal resistance to Azithromycin, Cefotaxime, and Ceftriaxone (9.09%). None were resistant to Meropenem, Cefuroxime, or Amoxicillin. All cadmium-resistant isolates (100%) displayed Multi-Antibiotic Resistance (MAR) to three or more antibiotics, highlighting significant public health concerns.

For chromium-resistant isolates, predominant resistance was observed to Tetracycline (83.33%), with Ampicillin and Nalidixic Acid each showing 50% resistance. Notable resistance was also seen to Ceftazidime (41.67%) and, to a lesser extent, to Ciprofloxacin, Cefotaxime, and Ceftriaxone (25%). Resistance rates for Gentamycin, Levofloxacin, Cefixime, Chloramphenicol, and Azithromycin were 16%, while resistance to Amoxicillin was 8%. No resistance was found to Cefuroxime or Meropenem. The MAR index for chromium-resistant isolates ranged from 0.187 to 0.437, indicating differing degrees of antibiotic resistance among the isolates. **(Fig 2)**

**Fig 2:**
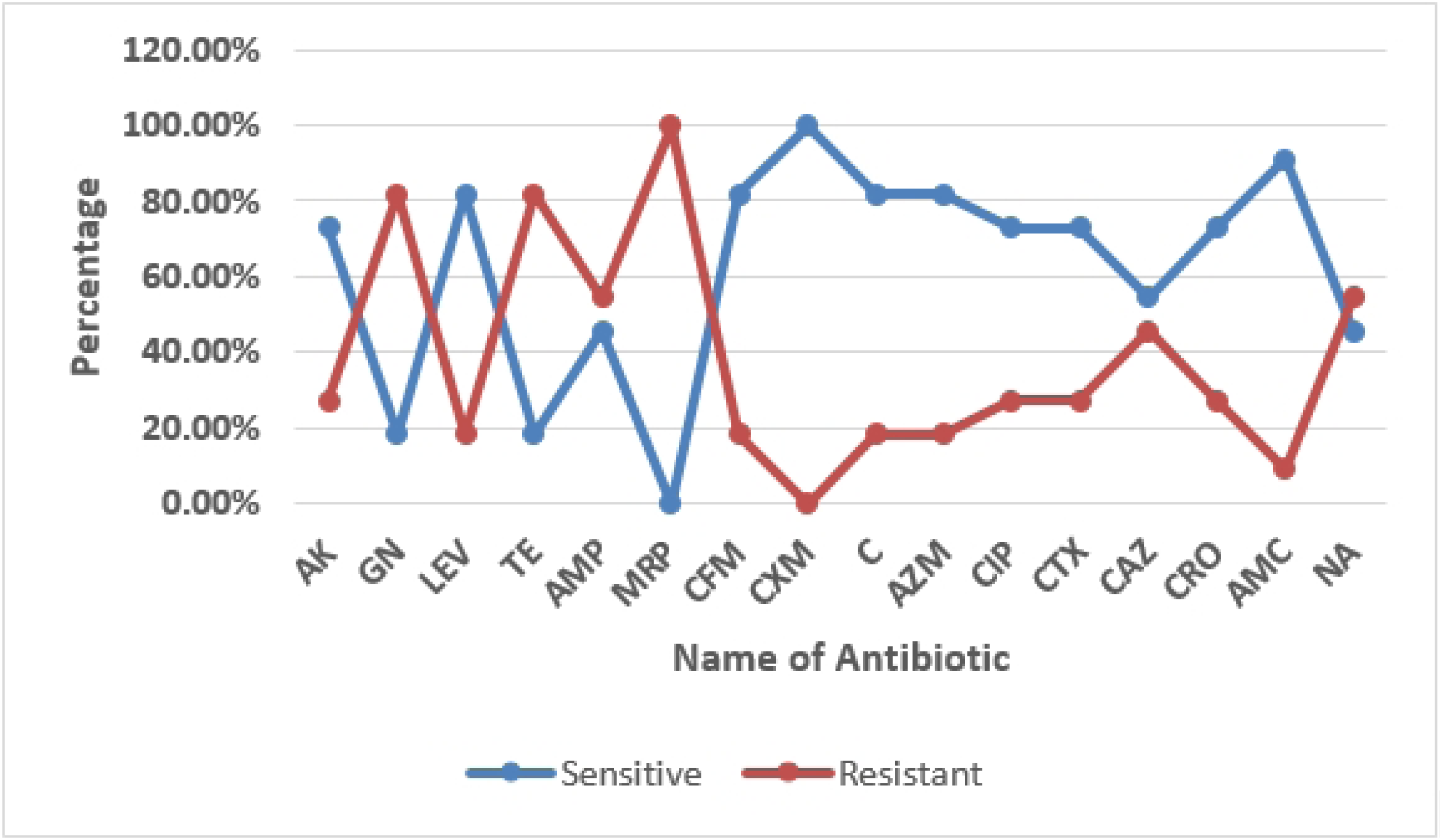
Antibiotic susceptibility pattern of chromium-resistant isolates against 16 different antibiotics. The highest percentage of resistance was found against Tetracycline (83.33%) and the lowest for Amoxicillin (8%).

### Molecular detection of heavy metal resistance genes

A total of twenty-three cadmium-resistant isolates were analyzed for cadmium-resistant genetic determinants. Isolates were targeted for the presence of *czc* for cadmium resistance. PCR of the isolates using a primer specific for cadmium resistance gene (*czc*) determined the presence of *czc* gene in six isolates (26%, 6/23).

PCR amplification of the metal-resistant genotype revealed that out of twenty-eight chromium-resistant isolates, five (18%, 5/28) contained chromium-reductase *chR* gene**. (Fig 3)**

**Fig 3:**
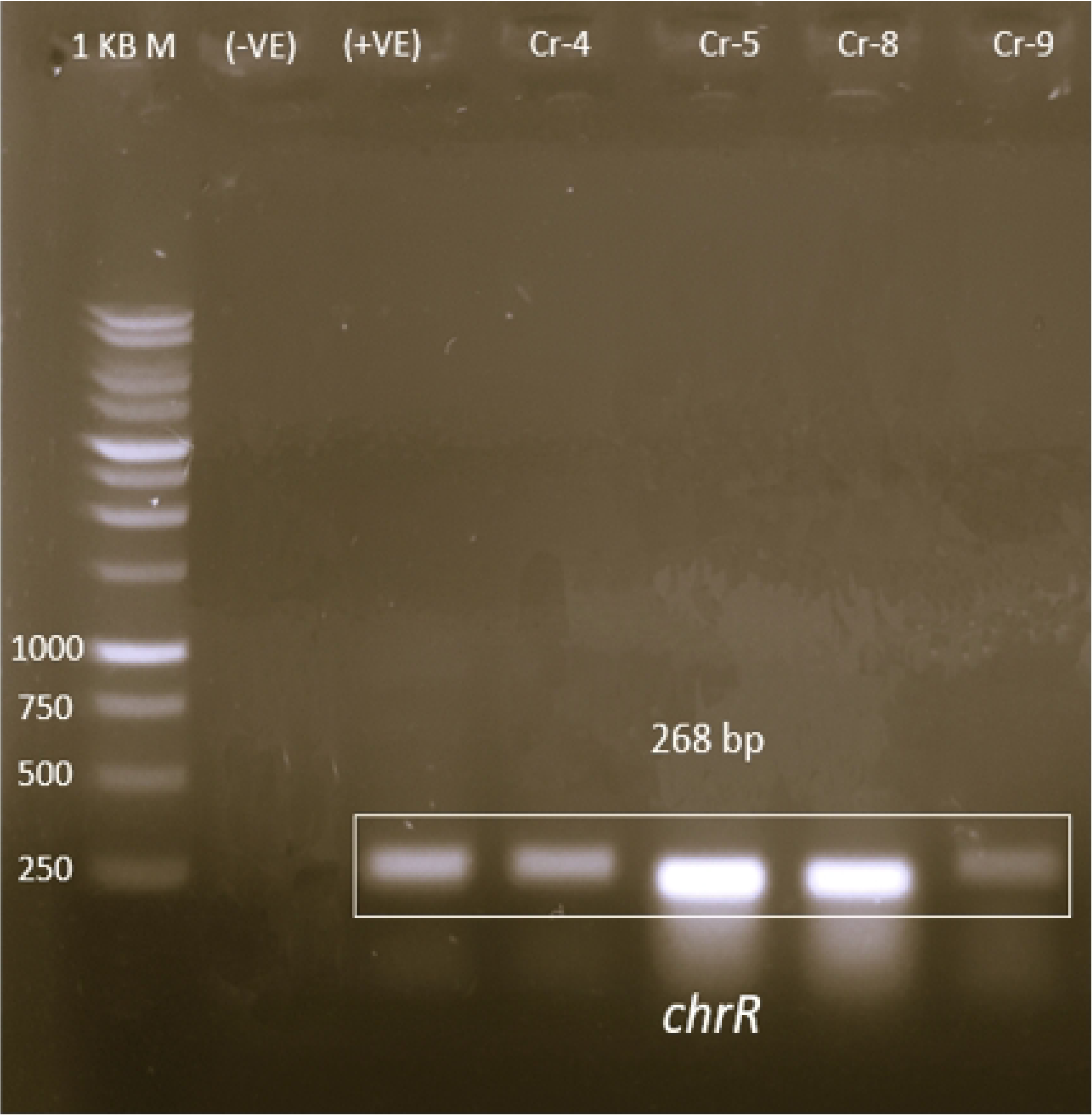
Heavy metal resistant gene *chrR* analysis for chromium resistant isolates. Lane 1:1 kb DNA Marker, Lane 2: Blank negative control, Lane 3:Positive control for *chrR*, Lane 4,5,6,7:Chromium resistant isolate positive for *chrR*.

### Antibiotic resistance determinants among metal resistance isolates

Screening five chromium-resistant isolates containing the *chrR* gene for resistance determinants against a range of antibiotics reveals a notable presence of genes conferring resistance to beta-lactam antibiotics and colistin. Specifically, the detection of *bla-TEM* **(Fig 4)** and *bla-NDM* in two isolates each (40%, 2/5) highlights the presence of extended-spectrum beta-lactamase (ESBL) and carbapenemase production, respectively. Furthermore, identifying the *mcr-2* gene in two isolates (40%, 2/5) is particularly concerning as it confers resistance to colistin, a last-resort antibiotic used to treat multi-drug resistant bacterial infections. The Chi-square test (p value = 0.026) also revealed a significant statistical association between heavy metal resistance genes and antibiotic resistance genes.

**Fig 4:**
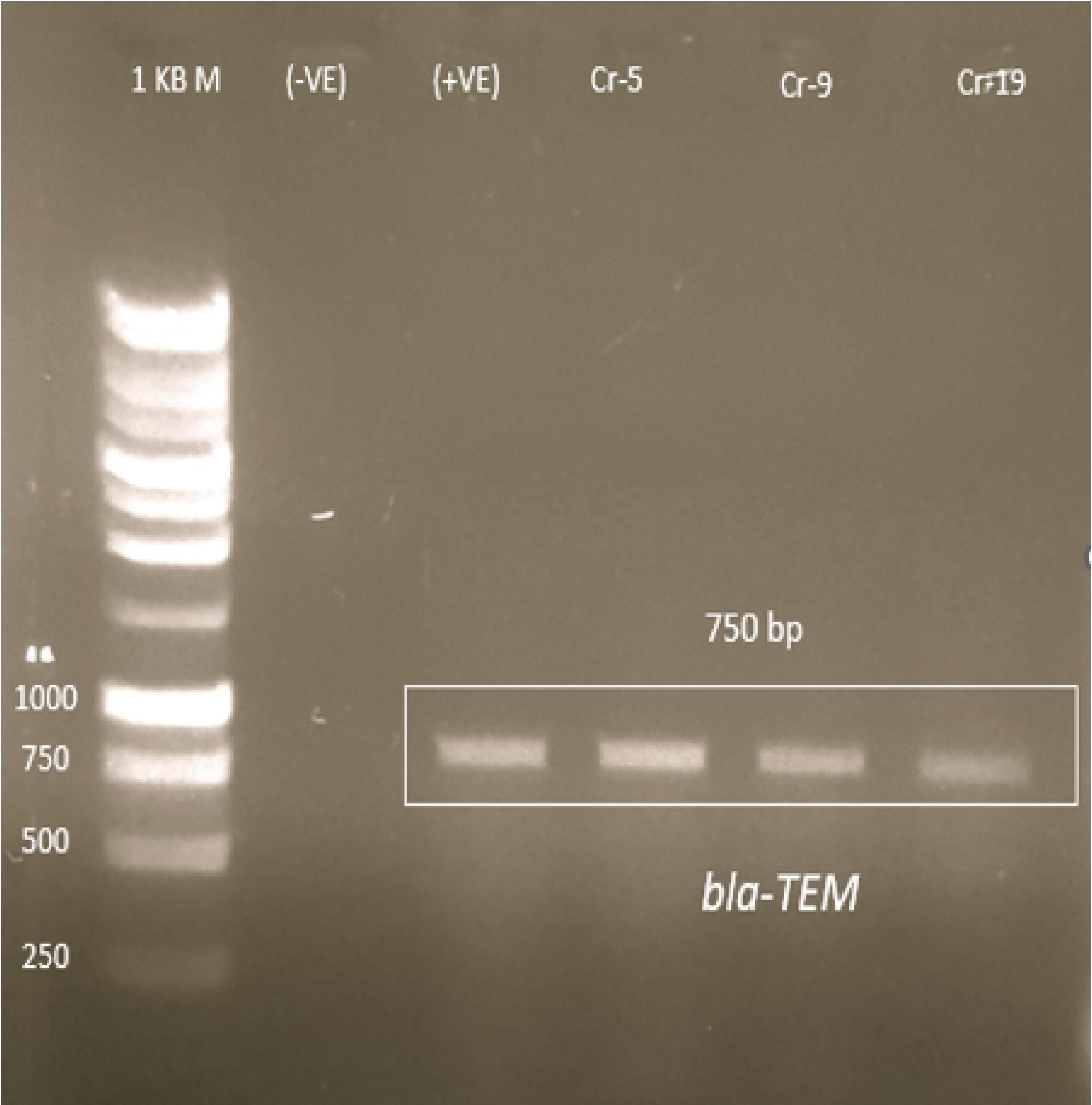
Antibiotic-resistant gene analysis among heavy metal-resistant isolates containing the metal-resistant gene. Lane 1:1 kb DNA Marker, Lane 2: Blank negative control (A) Lane 3: Positive control for *bla-TEM*, Lane 4,5: Chromium resistant isolates: Cr-5, Cr-9 containing *chrR* gene.

### Integron Analysis

Integrons were investigated in six isolates resistant to cadmium, which contained the *czc* gene, and five isolates resistant to chromium, containing the *chrA*gene. It was observed that two of the cadmium-resistant isolates with the *czc* gene harboured class 1 integrons (33.33%, 2 out of 6). All five chromium-resistant isolates with the *chrR* gene contained class 1 integrons (100%, 5 out of 5) **(Fig 5)**. The phenotypic resistance patterns and the corresponding genes detected in metal-resistant isolates from various poultry farms are summarized in **Table 2**.

**Fig 5:**
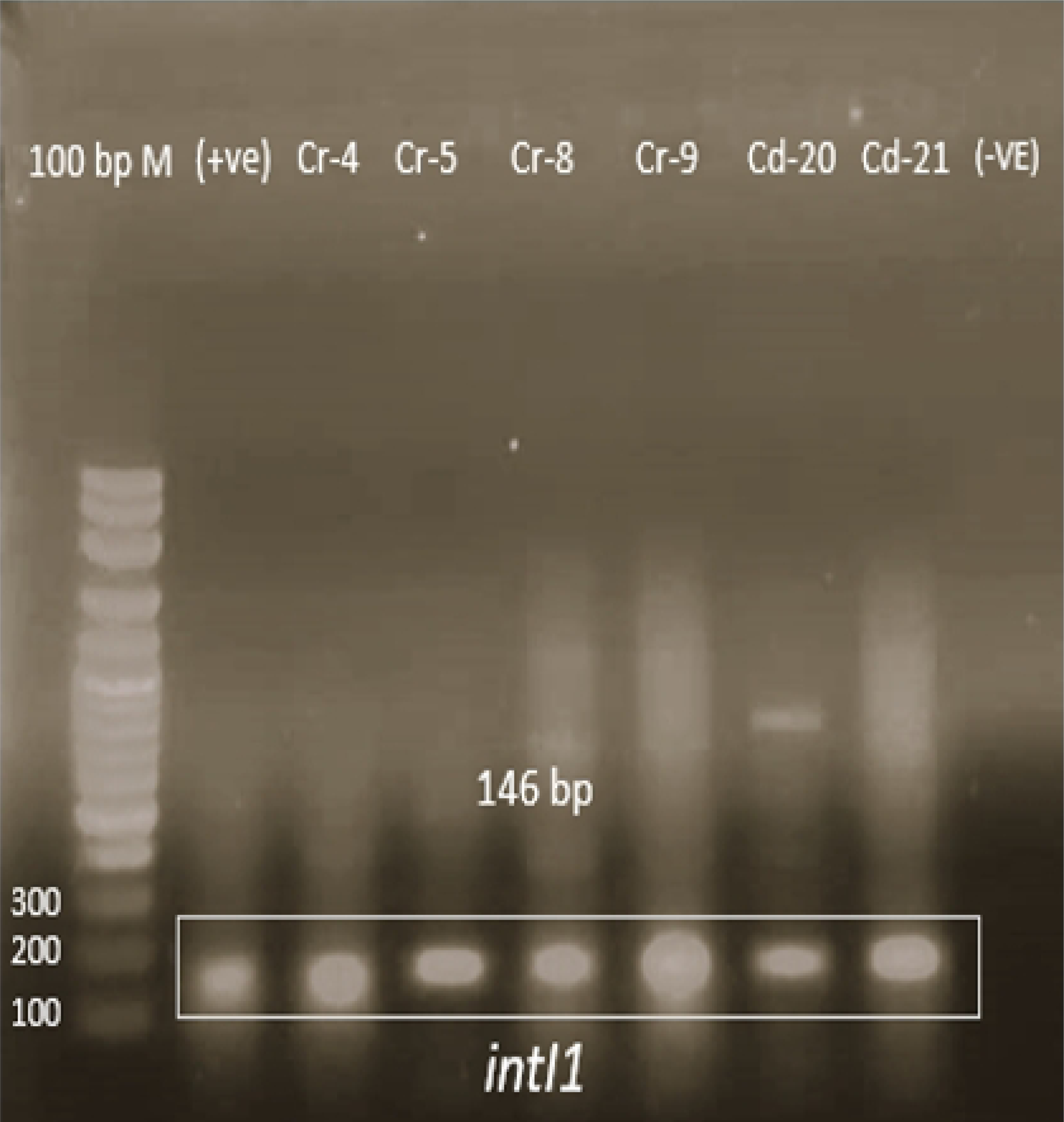
Detection of class 1 Inegron. Lane 1:100 bp DNA Marker, Lane 2: Positive control for *int 1* gene, Lane 3-6: Chromium resistant isolates:Cr-4, Cr-5, Cr-8, Cr-9 containing *chrR* gene.Lane7, 8: Cadmium resistant isolates:Cd-20, Cd-21 containing *czc* gene.

**Table 2:** Relative comparison among metal-resistant isolates from various poultry farms.

| Isolate ID | Sources | Tolerance to heavy metal | Multimetal resistance Pattern<sup>c</sup> |  |  |  | Antibiotic susceptibility test<sup>d</sup> | MAR Index | Heavy metal resistance gene | Antibiotic resistance gene | Integrase gene |
| --- | --- | --- | --- | --- | --- | --- | --- | --- | --- | --- | --- |
|  |  |  | Ni | Pb | Cad | Chr |  |  |  |  |  |
| Cd-11<sup>a</sup> | Droppings | 1000 µg/ml Cadmium | - | + | + | - | NA, GEN, LEV, CIP, CFM, C, CAZ | 0.46 | *czc* | NT<sup>e</sup> | NF<sup>f</sup> |
| Cd-12 | Droppings | 1000 µg/ml Cadmium | - | - | + | - | CFM, NA, CIP, LEV, C, CAZ | 0.37 | *czc* | NT | NF |
| Cd-16 | Droppings | 200 µg/ml Cadmium | + | + | + | - | CIP, NA, CFM, CAZ | 0.25 | *czc* | NT | NF |
| Cd-17 | Water | 100 µg/ml Cadmium | + | + | + | - | AK, CFM, AMC | 0.18 | *czc* | NT | NF |
| Cd-20 | Water | 100 µg/ml Cadmium | - | - | + | - | CFM, AK, AMC | 0.18 | *czc* | NT | *int 1* |
| Cd-21 | Water | 100 µg/ml Cadmium | + | + | + | - | CFM, AK, AMC | 0.18 | *czc* | NT | *int 1* |
| Cr-1<sup>b</sup> | Droppings | 1600 µg/ml Chromium | - | - | - | + | AK, LEV, NA, TE, AMP, CIP, CAZ | 0.43 | *chrR* | NF | *int 1* |
| Cr-4 | Droppings | 1600 µg/ml Chromium | - | + | - | + | GN, NA, TE, AMP, CAZ, C, CTX | 0.43 | *chrR* | NF | *int 1* |
| Cr-5 | Droppings | 400 µg/ml Chromium | - | + | - | + | AK, NA, TE, AMP, CIP, CRO | 0.37 | *chrR* | *bla-tem* | *int 1* |
| Cr-8 | Droppings | 400 µg/ml Chromium | - | - | - | + | NA, TE, AMP, CIP | 0.25 | *chrR* | *mcr-2, bla-ndm* | *int 1* |
| Cr-9 | Droppings | 400 µg/ml Chromium | - | - | - | + | NA, TE, AMP, C, CAZ | 0.312 | *chrR* | *bla-tem, mcr-2, bla-ndm* | *int 1* |

<sup>a</sup> Cd-Cadmium Resistant Isolates; <sup>b</sup> Cr-Chromium Resistant Isolates ; <sup>c</sup> Ni-Nickel ,Pb-Lead, Cad-Cadmium, Chr- Chromium; <sup>d</sup> NA- Nalidixic Acid, GEN- Gentamicin, LEV- Levofloxacin, CIP-Ciprofloxacin, CFM- Cefixime, C- Chloramphenicol, CAZ-Ceftazidime , AK-Amikacin, AMC-Amoxicillin-Clavulanic Acid, TE- Tetracycline, CTX- Cefotaxime, CRO-Ceftriaxone ; <sup>e</sup>NT means Not Tested, <sup>f</sup>NF means Not Found

## Discussion

Prior research has highlighted the influence of metals in promoting antibiotic resistance genes (ARGs) within manure [37] or soils enriched with manure [14,38]. However, from our current understanding, there needs to be more investigation into the simultaneous presence of heavy metal and antibiotic resistance within bacteria from the various poultry farms of Bangladesh, which represent the country’s largest meat production sector [39]. Our study uncovers these poultry farms’ concurrent patterns of metal resistance elements (MRE), ARGs, and mobile genetic elements (MGEs). Historically, including a varied mix of antibiotics and metals in poultry feed was a widespread practice to enhance growth rates and immunity [40], a practice now outlawed in Bangladesh and globally. Moreover, the use of tannery solid waste in poultry feed and aquaculture has been banned within the country to minimize harmful impacts [41]. These regulatory efforts aim to reduce the spread of antibiotic resistance genes (ARGs) in the environment, promoting sustained improvements in public health.

Given the contemporary context of poultry farming in Bangladesh, this study aimed to undertake a comprehensive screening and molecular characterization of isolates resistant to heavy metals and antibiotics and evaluate the bioremediation capabilities of heavy metal-resistant strains derived from the poultry sector in Bangladesh.

Various environmental factors, which include temperature and pH, greatly influence the growth and diversity of microorganisms. Temperature primarily affects microbial species’ physiological functions and distribution among the abiotic factors. Furthermore, variation in these parameters may affect microbial survival and their response to antibiotic pressure [42, 43], suggesting these conditions may contribute to the high resistance observed. The temperatures and pH levels of the study samples were found to meet the limits established by the Department of Energy (DoE) standards for Bangladesh [44]. This finding is consistent with various other studies conducted previously [42].In our study, dissolved oxygen (DO) levels fluctuated below 5 mg/L, slightly below the recommended level of 4.8 to 8 mg/L as per DoE [44]. Previous research has documented DO levels in poultry waste effluent ranging from 3.18 to 5.48 mg/L [45], indicating persistent environmental conditions that may enhance microbial resistance.

The biochemical tests in this study identified four bacterial genera: *Enterococcus spp., Escherichia coli, Staphylococcus spp.,*and *Bacillus spp*. The detection of *E.coli*, a key indicator of fecal contamination, in both poultry droppings and water samples highlights the need for better water quality monitoring in poultry farming areas. Previous studies have also noted E. coli and Staphylococcus spp., recognizing poultry as a reservoir for pathogenic strains [46]. Additionally, the identification of *Bacillus spp.* and *Enterococcus spp.*, both of which are known to carry antibiotic-resistant genes and may cause opportunistic infections, emphasizes the necessity for better waste management practices. This would help mitigate the risk of waterborne bacterial transmission. [47].

The study reveals significant resistance levels in bacteria from poultry environments against heavy metals like cadmium and chromium. Cadmium-resistant isolates showed MICs from 200 μg/ml to 1000 μg/ml, with 30% tolerating levels above 1000 μg/ml. Chromium-resistant isolates fully tolerated 200 μg/ml, with 21% resisting levels above 1600 μg/ml. These findings indicate a notable increase in resistance compared to previous reports for *E. coli* from poultry environments, where MICs were 25–400 μg/ml for cadmium and 400–800 μg/ml for chromium [48]. The widespread application of heavy metals in animal production and other sectors, including industry and agriculture, has led to their frequent presence across various settings. These metals are known to promote a selective advantage for bacteria that develop resistance, further highlighting the implications of their use in agriculture and other industries [4,49].

The results of this study show that nearly half of the chromium-resistant isolates exhibited co-tolerance to nickel and lead. In contrast, over half of the cadmium-resistant isolates were resistant to at least two other heavy metals. This indicates a substantial level of multiple heavy metal resistance, suggesting these isolates can survive in contaminated environments, posing potential environmental and health risks. Our study found lower resistance levels compared to previous research, such as the study on *E. coli* isolates from chicken neck skin and sheep cecum, which reported even higher resistance to nickel and zinc. [50]. The broader resistance spectrum to Chromium and Cadmium seen in our study, compared to Nickel and Zinc, highlights the urgency of addressing heavy metal resistance, particularly related to heavy metals in feed, to prevent the further spread of resistant strains. These findings underscore the need for enhanced monitoring and management strategies to address the growing issue of heavy metal resistance in poultry environments.

This study’s high incidence of multidrug resistance (MDR) raises serious concerns, mainly as all cadmium-resistant isolates were fully resistant to cefixime and ceftazidime. Chromium-resistant isolates also demonstrated alarming resistance levels, with 90% resistant to tetracycline and over half to ampicillin and nalidixic acid. This result suggests that exposure to heavy metals like cadmium and chromium may significantly contribute to the proliferation of MDR bacteria. Previous research on metal-resistant *E. coliform* chickens has documented similarly concerning resistance profiles, particularly against ampicillin, streptomycin, tetracycline, and ciprofloxacin, emphasizing the widespread issue of antibiotic resistance in metal-exposed environments. Additionally, studies on chromium-resistant *Salmonella strains* have shown significant resistance to cefotaxime, further illustrating the variability in resistance mechanisms across bacterial species and their responses to different heavy metals [51]. The juxtaposition of these studies illuminates the intricate relationship between heavy metal exposure and the development of AR. Notably, the impact of heavy metals on antibiotic resistance genes (ARGs) was highlighted as potentially more significant than antibiotics themselves in some contexts. This result suggests that the selective pressure exerted by heavy metals in the environment might favour the survival of metal-resistant bacteria and facilitate the co-selection of ARGs, thus accelerating the spread of MDR.

The utilization of heavy metals such as Cu, Zn, Mn, Cr and Ca in livestock farming is widespread due to their benefits in promoting growth and preventing animal diseases. However, this intensive application leads to significantly elevated levels of these metals in livestock manure and the surrounding environment, posing potential risks of heavy metal contamination [10, 52,53]. Our study focused on the resistance mechanisms against heavy metals such as cadmium. We found that the *czc* gene, which confers resistance to cadmium, was detected in one-fourth of cadmium-resistant isolates. The czc system, known for its role in resisting Cd^2+^, Zn^2+^, and Co^2+^ operates through an active efflux pump complex composed of CzcA, CzcB, and CzcC proteins, functioning as a cation-proton antiporter [54,55]. Comparatively, previous research on *Aliarcobacter butzleri* strains from German water poultry identified the presence of *czcA*and *czcB* genes, responsible for encoding cobalt-zinc-cadmium resistance proteins, were detected in 22% and 17% of strains [56]. This comparison highlights the critical role of specific genes and efflux systems in conferring resistance to heavy metals across different bacterial species. The prevalence of such resistance mechanisms underscores the environmental and public health challenges posed by the widespread use of heavy metals in poultry and their consequent accumulation in the environment. Addressing these challenges requires a comprehensive understanding of resistance mechanisms and practical strategies to mitigate the impact of heavy metal contamination.

Chromate Resistance Determinants (CRDs), including the *chrA* and *chrR* gene, are widespread across various life domains and are crucial for chromate resistance [57, 58]. The role of the chrR gene in the bacterial chromate resistance mechanism is well-established, with chromium reductase, which reduces Cr(VI) to Cr(III), being found in chromosomal DNA, plasmids, or both [59,60]. Our study identified the *chrR*gene, associated with chromium reduction, in a subset of the chromium-resistant isolates, indicating its notable presence in specific bacterial communities. In contrast, earlier research found a 100% occurrence of the *chrR* gene in chromium-resistant bacteria from poultry droppings, suggesting a higher prevalence in that specific environment [61]. The variation in prevalence rates across different studies and environments underscores the dynamic nature of resistance gene distribution, influenced by factors such as antibiotic use, heavy metal exposure, and agricultural practices. The non-degradable nature of heavy metals, such as chromium, poses a long-term selection pressure on microbial populations. This environmental challenge drives the development and maintenance of heavy metal resistance genes (HMRGs), including *chrA*, contributing to the persistence and spread of resistance mechanisms over time [62]. The enduring presence of heavy metals in the environment underscores the need for sustainable practices that minimize heavy metal pollution and its role in promoting resistance.

The findings from the present study provide a compelling illustration of the interconnectedness between heavy metal resistance genes (HMRGs) and antibiotic resistance genes (ARGs), particularly within the context of chromium resistance mediated by the *chrR* gene. Our study identified notable antibiotic resistance genes such as *bla-TEM*, *bla-NDM,*and *mcr-2* in chromium-resistant isolates harbouring the *chrR* gene, suggesting a significant overlap between resistance mechanisms against heavy metals and antibiotics. The significant statistical association (p-value = 0.026) between heavy metal resistance genes and antibiotic resistance genes implies that resistance to one may promote or co-select for resistance to the other. This is likely due to the shared genetic elements or environmental pressures that select for both types of resistance. This co-occurrence is indicative of a broader resistance landscape where the presence of HMRGs like *chrR* can be closely linked to ARGs conferring resistance to various antibiotic classes. This observation supports previous research showing associations between HMRGs (e.g., *arsB* and *chrA*) and ARGs responsible for β-lactam resistance (*blaCTXM, blaTEM, blaSHV*) in organisms like Salmonella [48]. Such associations have been statistically significant, suggesting a robust correlation between the presence of heavy metal resistance and the propensity for antibiotic resistance. This correlation is likely due to the co-localization of HMRGs and ARGs on the same mobile genetic elements (MGEs), such as plasmids and integrons, facilitating their co-selection and dissemination among bacterial populations [62–64]. The implications of these findings are significant, suggesting that strategies to combat antibiotic resistance must consider the environmental context, including the presence and impact of heavy metals. Efforts to reduce the discharge of heavy metals into the environment could mitigate the selection pressure for both HMRGs and ARGs, potentially slowing the spread of resistance. Moreover, understanding the genetic and ecological dynamics that facilitate the co-occurrence of HMRGs and ARGs could inform the development of targeted interventions to disrupt the spread of resistance genes among bacterial populations.

Mobile genetic elements, including transposons, integrons, and insertion sequences, are paramount in the evolutionary dynamics of antibiotic resistance genes (ARGs) within microbiomes. These elements facilitate horizontal gene transfer among microorganisms, enabling the dissemination of ARGs in diverse environments [15,16,18]. Class 1 integrase (intI-1) genes, for instance, have been identified as prevalent in bacteria present in poultry litter, regardless of antibiotic usage [65]. Furthermore, class 1 integrons are frequently detected among Enterobacteriaceae, with the *chrA* gene often found downstream of the 3ʹ-conserved segment (CS) of the class 1 integron in plasmids. This arrangement suggests a potential mechanism for co-resistance to chromate and antibiotics [66, 67]. In our study, class 1 integrons were observed in cadmium-resistant isolates with the *czc* gene and chromium-resistant isolates with the *chrR* gene. In a previous study, a similar genetic structure, including *intI1–sul1–chrA,* has been reported in Salmonella genomic island 1 (SGI1), indicating a possible genomic island formation. This suggests that chromium-resistant genes and class 1 integron may constitute a genomic island that spreads collectively [67]. These findings illuminate the interplay between mobile genetic elements, ARGs, and resistance to heavy metals and antibiotics. Further research into the dynamics of mobile genetic elements and their influence on ARG and HMRG dissemination is warranted to develop targeted interventions for combating antibiotic and metal resistance.

Overall, this research marks the first comprehensive investigation in Bangladesh to explore the co-occurrence of heavy metal and antibiotic resistance in bacteria from poultry farms. The study’s findings highlight the urgent need for stricter regulation of antibiotic, heavy metal, and disinfectant usage in the poultry industry to curb the spread of resistance. The study reinforces the importance of implementing One Health approaches to address the ongoing antimicrobial and heavy metal resistance spread, ensuring a holistic strategy that integrates human, animal, and environmental health.

## Conflict of Interest

The authors have no financial conflicts of interest to declare.

## Notes

### Competing Interest Statement

The authors have declared no competing interest.

